# Classification of the gifted natural product producer *Streptomyces roseofaciens* sp. nov. by polyphasic taxonomy

**DOI:** 10.1101/310888

**Authors:** Lizah T. van der Aart, Imen Nouinoui, Alexander Kloosterman, José Mariano Ingual, Joost Willemse, Michael Goodfellow, Gilles P. van Wezel

## Abstract

A novel verticillate strain of streptomycetes, *Streptomyces* strain MBT76^T^, was isolated from the QinLing mountains, which harbours more than 40 biosynthetic gene clusters for natural products. Here we present full taxonomic classification of strain MBT76^T^, and show that it has chemotaxonomic, genomic and morphological properties consistent with its classification in the genus *Streptomyces*. Strain MBT76^T^ is part of the cluster of Streptoverticillates, a group within the genus *Streptomyces* that has characteristic whorl-forming spores produced in chains along the lateral wall of the hyphae. Multi-locus sequence analysis based on five housekeeping gene alleles showed that MBT76^T^ is closely related to *Streptomyces hiroshimensis*. Average Nucleotide Identification (ANI) and Genome to Genome Distance Calculation (GGDC) of the genomes of strain MBT76^T^ and *S. hiroshimensis* separated them into distinct species. Strain MBT76^T^ represents a novel species of the genus *Streptomyces* for which we propose the name *Streptomyces roseofaciens* sp. nov. The type strain is MBT76^T^ (=NCCB 100637^T^ =DSM 106196^T^). The whole genome of MBT76^T^ has 7974 predicted open reading frames and a total genome size of 8.64 Mb. Further genomic analysis showed that verticillate streptomycetes lack the sporulation gene *ssgE*, and our data suggest that this is a useful genetic marker for the spore-chain morphology of the verticillates.

## INTRODUCTION

Actinomycetes are a major source of secondary metabolites, among which antimicrobial, antifungal and anticancer compounds (1). Of these, the streptomycetes are an especially rich source of clinical secondary metabolites, and produce over half of all clinically used antibiotics (2). One of the challenges of antibiotic discovery is that the potential for secondary metabolite expression of *Streptomyces* is difficult to determine. Even the model organism *Streptomyces coelicolor,* which has been a topic of research of over 30 years, produces unexpected secondary metabolites under novel conditions (3). Classification of prokaryotes is a rapidly changing field, being driven by technological advances like the current increase in genome sequence information. Even though the field is continuously leaning more towards the use of molecular techniques, phenotypic data needs to be collected to draw conclusions on the novelty of a species, for this reason we use a polyphasic approach (4–7). Classification of prokaryotes is typically based on genomic and phenotypic data, *i.e*. on polyphasic taxonomy (5, 6, 8). The impact of 16S rRNA gene sequence and DNA:DNA relatedness values is particularly high in terms of delineating taxa at the rank of species (4, 9), and has led to strongly improved classification of taxa belonging to the phylum Actinobacteria (7). Still, the resolution offered by 16S rRNA sequences is not sufficient for the recognition of new taxa. The SsgA-like proteins (SALPs) are very good additional markers for the accurate classification of Actinobacteria, and relate closely to morphological characteristics (10). The SALPs are unique to morphologically complex actinobacteria (11), and orchestrate aspects of peptidoglycan synthesis and remodelling, including cell division and spore maturation (12, 13). The archetypal SALP is SsgB, which initiates sporulation-specific cell division (13). SsgB shows extremely high conservation (maximum of one aa change) within a genus, while there is high diversity even between closely related genera, making it a good marker to help classify genera within the Actinobacteria (10). Members of the genus *Kitasatospora* have an SsgB orthologue that differs from that of streptomycetes in 4 positions, which is a distance that is sufficient to separate the two genera (14).

The position of the genus *Streptoverticillium* (15) has been a subject of debate for decades, and it is now generally accepted based on 16S rRNA phylogeny that members of this group of verticillate, or whorl-forming, *Streptomycetaceae* in fact belong to the genus *Streptomyces* (10, 16). Exceptionally, while the aerial hyphae of most streptomycetes develop into a single chain of spores at the apex, the verticillates produce small chains of spores at multiple sides perpendicular to the aerial hyphae. The MBT-collection of Actinobacteria isolated from soil from the QinLing mountains in China display wide phylogenetic and chemical diversity (17). We recently identified a novel member of the verticillate streptomycetes, namely *Streptomyces* species MBT76^T^. A screen against the so-called ESKAPE pathogens (18) followed by genome and natural product mining showed that *Streptomyces* sp. MBT76^T^ is a rich source of natural products, including antibiotics (19). Analysis of the genome using AntiSMASH (20) identified 44 putative biosynthetic gene clusters (BGCs) for the secondary metabolites. Further investigation of the natural products produced by MBT76^T^ using NMR-based metabolomics (21) identified a range of bioactive compounds, including isocoumarins, flavonoids, phenylpropanoids, siderophores and naphtaquinones (22–25).

In this work we wish to establish strain MBT76^T^ as a novel species of *Streptomyces*. MBT76^T^ was originally isolated from the QinLing mountains and has been the subject of several metabolomic studies. Polyphasic taxonomy shows that MBT76^T^ is a novel verticillate *Streptomyces* species for which we propose the name *Streptomyces roseofaciens* sp. nov., the name reflecting its production of a red/pink pigmented compound.

## MATERIAL METHODS

### Media and growth conditions, strains

*Streptomyces* sp. MBT76^T^ was isolated from a soil sample collected from Shandi Village, the Qinling mountains, Xi’an, Shaanxi Province, China: (34°03’28.1”N, 109° 22’39.0”E) height 660 m (26). MBT76^T^ is part of the culture collection at Molecular Biotechnology, IBL, Leiden University. The reference strain, *Streptomyces hiroshimensis* DSM 40037 was obtained from the DSMZ collection. The strains were maintained by sub-culturing on ISP-2 and a spore-stock is frozen in glycerol at -80 degrees. Biomass for biochemical tests was harvested from solid ISP-2 medium and freeze-dried.

### Phylogenetic analysis

The complete 16S rRNA gene sequence (1,416 nucleotides [nt]), isolated from the genome sequence of *Streptomyces* sp. MBT76^T^ (Genbank accession number: LNBE00000000.1), was submitted to the EzTaxon-e server (http://eztaxone.ezbiocloud.net/; (27, 28) and aligned with corresponding 16S rRNA gene sequences of the type strains of the most closely related *Streptomyces* species using CLUSTALW version 1.8 (29). Phylogenetic trees were generated from the aligned sequences using the maximum-likelihood (30), maximum-parsimony (31) and neighbour-joining (32) algorithms drawn from the MEGA 5 and PHYML software packages (33, 34); an evolutionary distance matrix for the neighbour-joining analysis was prepared using the Jukes and Cantor model (35). The topology of the inferred evolutionary trees was evaluated by bootstrap analyses (36) based on 1,000 resamplings of the maximum-likelihood using MEGA 5 software. The root positions of unrooted trees were estimated using the sequence of *Kitasatospora setae* KM 6054^T^.

### Multilocus Sequence Analysis

Multilocus sequence analysis was based on the method of Labeda (37). The sequences of *atpD* (ATP synthase F1, β-subunit), *gyrB* (DNA gyrase B subunit), *recA* (recombinase A), *rpoB* (RNA polymerase β-subunit) and *trpB* (tryptophan synthase, β-subunit) were extracted from the full genome sequence of strain MBT76^T^. The sequences of the loci for each strain were concatenated head to tail and exported in FASTA format, providing a dataset of 33 strains and 2351 positions. Sequences were aligned using MUSCLE (38) and phylogenetic relationships constructed in MEGA 5.2 (33) using maximum-likelihood based on the General Time Reversible model (39). The phylogenetic relationships of the strains were also determined using maximum-parsimony and neighbour-joining analyses. MLSA evolutionary distances were determined using MEGA 5.2 to calculate the Kimura 2-parameter distance (40).

### Whole Genome analysis

The average nucleotide identity (ANI) between the genomes of MBT76 (GenBank Accession number: NZ_LNBE01000001.1) and *S. hiroshimensis* (GenBank Accession number: NZ_JOFL01000001.1) was determined using the OrthoANIu algorithm available as an online tool on EZbiotaxon (41). The digital DNA–DNA hybridization (dDDH) values between the genomes were calculated using the genome-to-genome distance calculator, GGDC 2.0 available at http://ggdc.dsmz.de. For dDDH, a cut-off value of 70% is used. (42). The ANI was calculated using the ANI calculator on EzBiocloud using the orthoANIu algorithm (43). For ANI, a general cut-off value of 95-96% was used.

### Sequence alignment and phylogenomic analysis

To find all SsgA-like proteins (SALPs) for the strains of interest, refseq annotated protein files were downloaded from NCBI of three verticillate strains of which a full genome sequence was available: *S. hiroshimensis* (NZ_JOFL01000001.1), *S. cinnamoneus* (NZ_MOEP01000440.1) and *S. mobaraensis* (NZ_AORZ01000001.1). *S. coelicolor* (NC_003888.3), *S. griseus* (NC_010572.1) and *S. lividans* TK24 (NZ_GG657756.1) were added as reference strains. All genes for SALPs were obtained from the genome sequences of these strains by a BLAST search with low cut-off (e-value 10^-5^) using the SALPs from *S. coelicolor* as the queries. To verify that all hits found were true SALPs, a second BLAST search was performed, using the output hits to interrogate the genome sequence of *S. coelicolor* M145. All hits whose reciprocal best hits were again SALPs were used for further phylogenetic analysis. Positive hits were then aligned using MUSCLE (38). Phylogenetic trees were generated using maximum-likelihood algorithms with default parameters as implemented in MEGA version 5 (33) The tree reliability was estimated by bootstrapping with 1,000 replicates (36). Secondary metabolite biosynthetic gene clusters were assessed using the antiSMASH 4.0 server (44).

### Chemotaxonomy and morphology

MBT76^T^ was examined for chemotaxonomic and morphological properties considered to be typical of the genus *Streptomyces* (45). The arrangement of aerial hyphae and spore chains were observed on oatmeal agar (ISP medium 3; (46)) after 14 days at 28°C. Spore chain morphology and spore surface ornamentation were detected by examining gold-coated, dehydrated specimens by Scanning Electron Microscopy (JEOL JSM-7600F instrument) (47). Cultural characteristics of the isolate were determined on ISP 1-7 media (46) following incubation at 28°C for 14 days. Standard protocols were used to detect the isomers of diaminopimelic acid (48), menaquinones and polar lipids (49), and whole organism sugars (48). Cellular fatty acids were extracted, methylated and analysed by gas chromatography (Hewlett Packard, model 6890) following the recommended procedure of the Sherlock Microbial Identification System (50). The resultant fatty acid methyl esters were identified and quantified using the MIDI ACTINO 1 database (version 6.10).

### Phenotypic tests

*Streptomyces* sp. MBT76^T^ and *S. hiroshimensis* DSM 40037^T^ were examined for biochemical, degradative and physiological properties, in duplicate, using media and methods described by Williams et al. (51) and known to be of value in the systematics of Streptomycetes (45). The enzyme profile of the strain was determined using API ZYM strips (BioMerieux) following the manufacturer’s instructions; a standard inoculum equivalent to 5.0 on the McFarland scale (52) was used to inoculate the API ZYM strips.

## RESULTS AND DISCUSSION

Strain MBT76^T^ has cultural and morphological properties typical of *Streptomyces* (45). *Streptomyces* sp. MBT76^T^ was isolated from a Qinling mountain soil sample (26) and produces many natural products that are activated in response to specific environmental cues (17). The metabolomic potential was assessed by NMR-metabolomics (24) and genomics (23). Strain MBT76^T^ shows verticillate sporulation with smooth spores (Figure 1). The strain grows moderately well on most ISP-media, and well on ISP2, 3 and 4 (Table 1), producing mostly pink pigments, characteristic of representatives of the red-pigmented verticillates (53).

**Figure 1:**
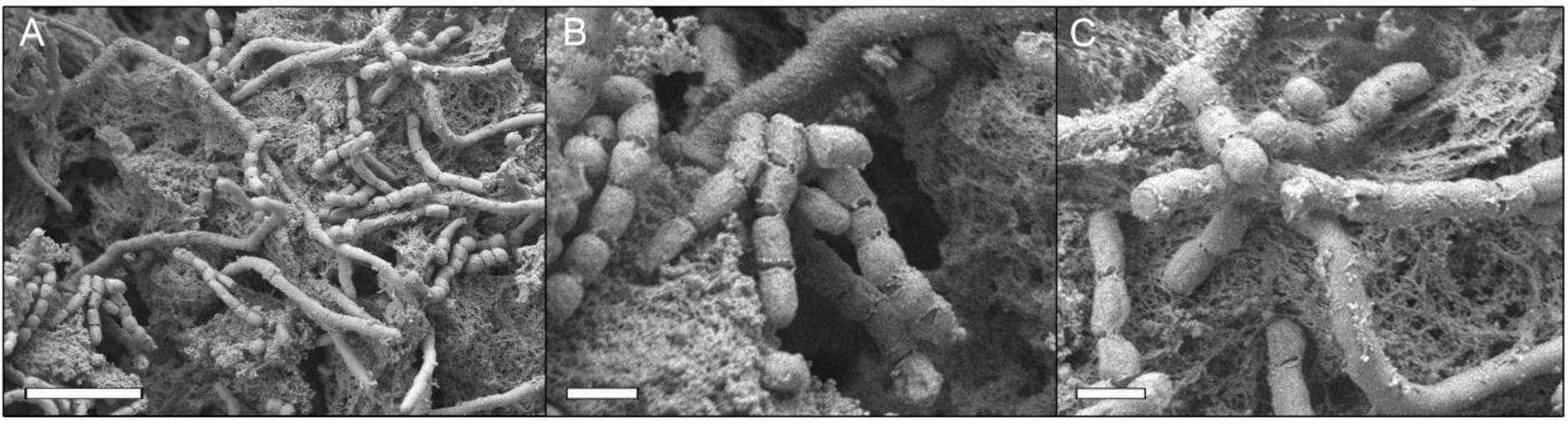
Scanning electron micrographs from a 14-day old culture of *Streptomyces* MBT76^T^ grown on ISP-3 agar plates. Images B and C are enlargements of image A. MBT76^T^ produces smooth, verticillate spores of +/− 1 μM in size. Bars: A: 5 μM; B, C: 1 μM.

### 16S rRNA and MLSA trees

When the 16S rRNA gene of strain MBT76 is compared with sequences of neighbouring *Streptomyces,* it was found to be closely related to *Streptomyces hiroshimensis* (99.37%), *Streptomyces mobaraensis* (99.24%) and *Streptomyces cinnamoneus* (99.17%). 16S rRNA gene sequence similarities with the remaining strains fell within the range of 99.10 to 98.13% (Figure 2). The results of the multi-locus sequence analysis (MLSA), based on five house-keeping genes head to tail, are shown in Figure 3. Isolate MBT76^T^ forms a well-supported clade with the type strain of *S. hiroshimensis^T^,* as the strain morphologically fits in a clade with red-pigmented verticillate streptomycetes of which the morphology and pigmentation clusters very well with MLSA-based phylogeny (53–55). The MLSA evolutionary distances between isolate MBT76 and other verticillate strains in the same clade are shown in Table 3. Strain MBT76 showed an MLSA distance greater than 0.007 with all phylogenetically near species, supporting the proposal that this strain represents a new species (Table 2) (56).

**Figure 2.**
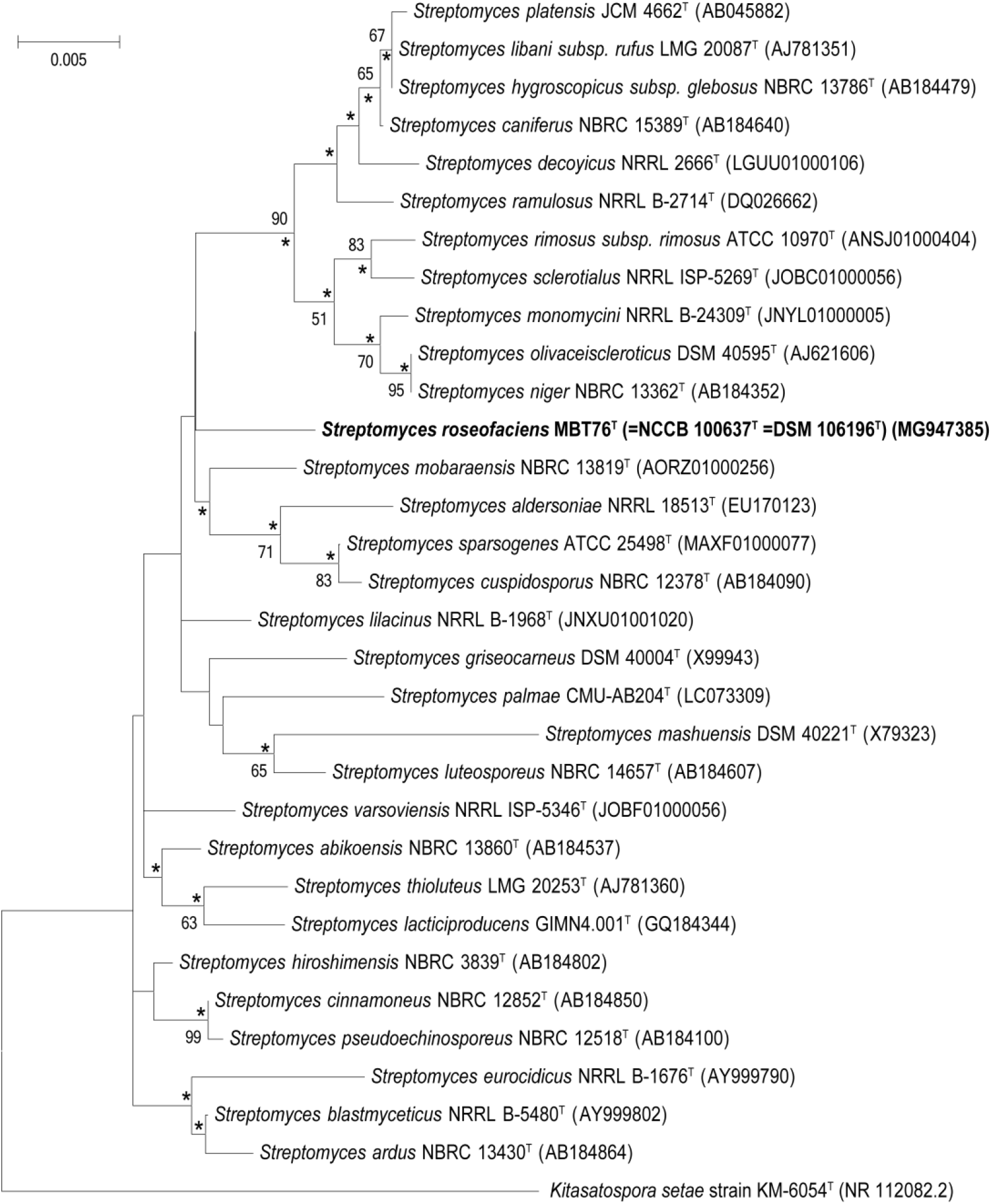
Maximum-likelihood phylogenetic tree (30) based on 16S rRNA gene sequences. The tree shows relationships between isolate MBT76 and the type strains of closely related *Streptomyces* species. Asterisks indicate branches of the tree that were also recovered using the neighbour-joining (32) and maximum-parsimony (62) tree-making algorithms. Numbers at the nodes indicate levels of bootstrap based on a maximum likelihood analysis of 1,000 sampled datasets, only values above 50% are given. The root position of the tree was determined using *Kitasatospora setae* KM-6054T. GenBank accession numbers are given in parentheses. Scale bar, 0.005 substitutions per nucleotide position.

**Figure 3.**
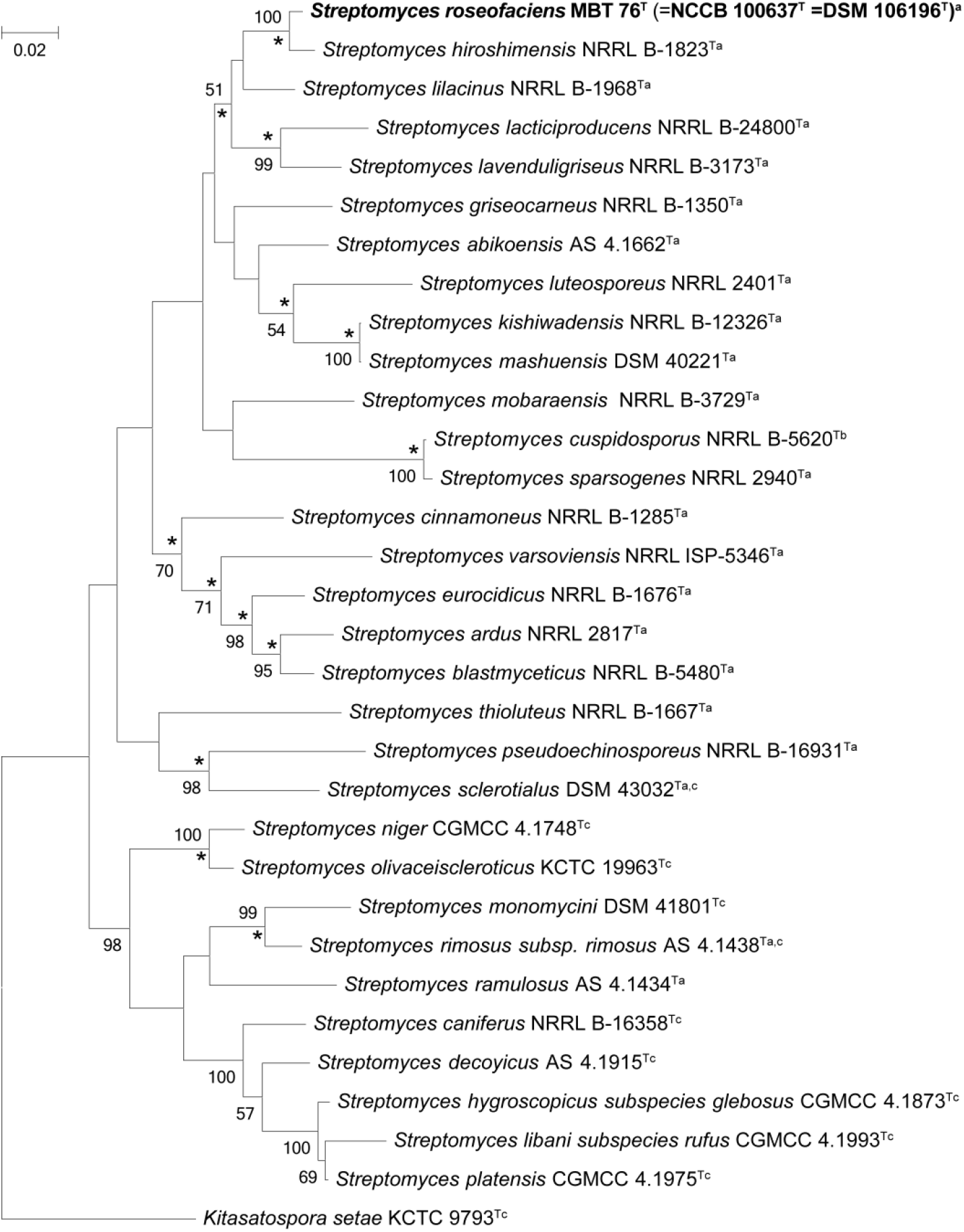
Phylogenetic tree based on MLSA analysis. *Streptomyces roseofaciens* (bold) is a part of a clade consisting of verticillate *Streptomyces*. Maximum-likelihood phylogenetic tree inferred from partial sequences of the house-keeping genes *atpD, gyrB, recA, rpoB* and *trpB* in MEGA 6.0. The analysis involved 33 nucleotide sequences with a total of 2351 positions in the final dataset. Neighbour-joining and maximum-parsimony models in Mega6.0 and conserved branches in all methods are marked with an *asterisk*. Percentages at the nodes represent levels of bootstrap support from 1,000 resampled datasets with values with less than 60% not shown. Streptomyces morphology: ^a^: verticillate spore chains. ^b^: not determined ^c^: *Streptomyces* with canonical (apical) spore chains.

To validate that strain MBT76^T^ and *S. hiroshimensis* are separate species, *in silico* DNA:DNA hybridisation (DDH) studies were performed using two different methods, both the genome-to-genome distance calculator (GGDC) (57–59) and Average Nucleotide Identity (ANI)(41). Comparisons with the genome of *Streptomyces* strain MBT76^T^ with *S. hiroshimensis^T^* yielded GGDC values of 28.40%± 2.3 %, confirming that the strain is genetically separated from the type strain of *S. hiroshimensis* (Table S1). The orthoANIu value of MBT76^T^ and *S. hiroshimensis* is 88.96 (Table S2). Both the GGDC and ANI values are well below the cut-off point for prokaryotic species delineation. The genomic DNA G+C composition of strain MBT76^T^ is 71.9%. Interestingly, *S. cinnamoneus* and *S. mobaraensis* are known to produce elfamycin-type antibiotics of the kirromycin class that target elongation factor EF-Tu, and the antiSMASH data of MBT76^T^ and *S. hiroshimensis* shows that these two strains are potential kirromycin producers as well (60, 61).

### Taxonomic classification based on SALP phylogeny

The developmental genes encoding members of the family of SsgA-like proteins (SALPs) serve as good predictive markers for morphological differences between Actinobacteria (10). Considering the obvious difference in spore-chain positioning between verticillates, which form small spore chains at multiple position along the lateral wall, and canonical streptomycetes that form long spore chains at the tips of the aerial hyphae, we compared the SALP proteins. Of these, SsgA, SsgB, SsgD, SsgE and SsgG are conserved within the streptomycetes, while various additional members are encountered with a strain- or clade-specific distribution. As verticillate streptomycetes we chose *Streptomyces* sp. MBT76^T^, *S. hiroshimensis, S. cinnamoneus,* and *S. mobaraensis,* and as non-verticillate *S. coelicolor, S. lividans* and *S. griseus* (Figure 4). The *ssgA* gene is a useful phylogenetic marker for the further subclassification of members of the genus *Streptomyces* (10). The SsgA-based branch confirms the close correlation between MBT76^T^ and *S. hiroshimensis*^T^.

**Figure 4.**
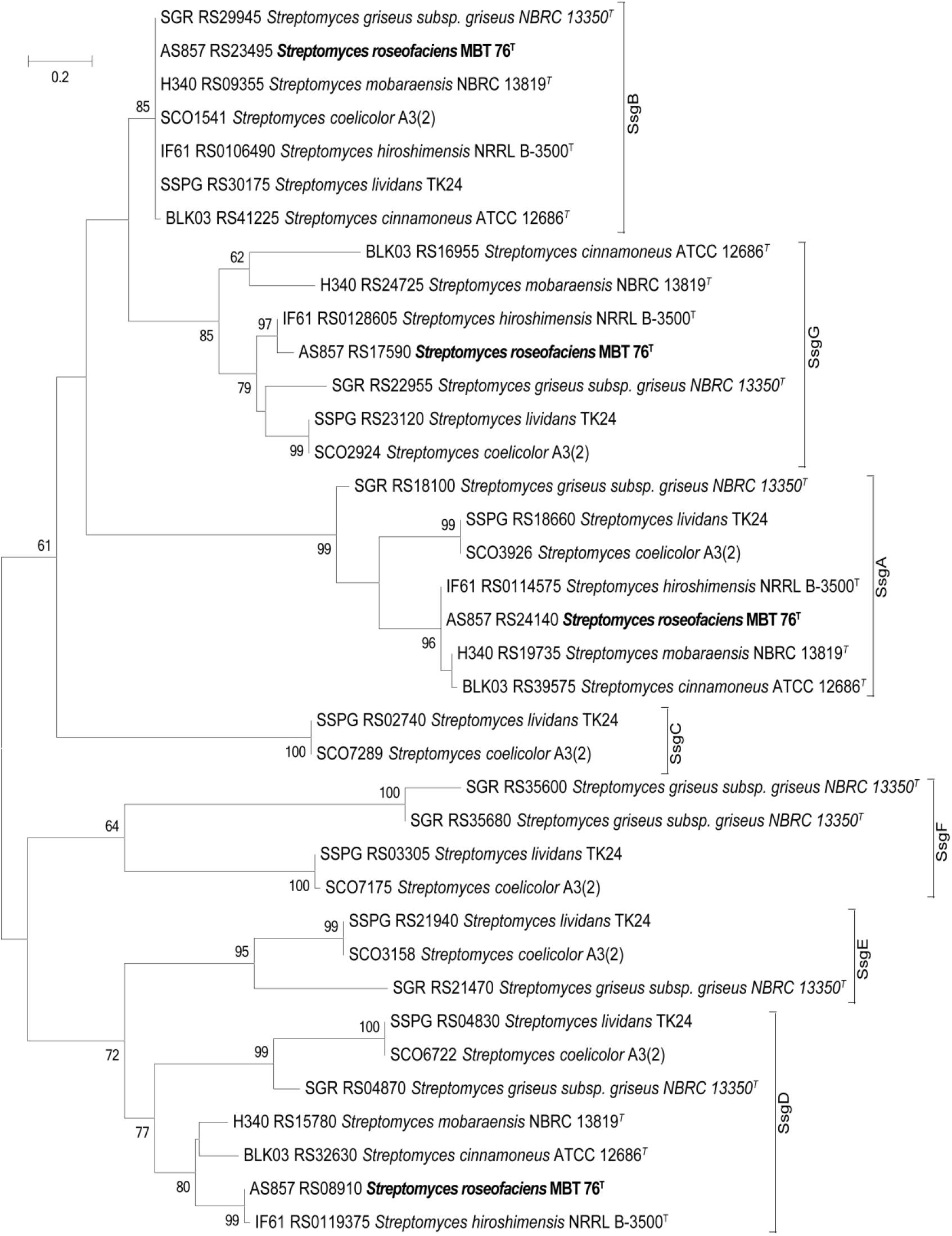
Phylogenetic tree based on SALP sequences. Maximum-likelihood phylogenetic tree of SALP protein sequences in *S. roseofaciens, S. hiroshimensis, S. cinnamoneus, S. mobaraensis, S. coelicolor, S. lividans* and *S. griseus*. The locus-ID of each SALP is indicated in the respective branch.

As mentioned above, the SsgB sequence of streptomycetes is invariable, except for amino acid residue 128. Streptomycetes that carry a threonine at position 128 (T128) sporulate in submerged cultures, while those carrying a glutamine (Q128) do not. Like all verticillates, strain MBT76 has an SsgB sequence that is consistent with classification within the genus *Streptomyces*. The specific variant is SsgB T128 variant, suggesting that the verticillates sporulate in submerged cultures. Preliminary experiments confirmed this ability for strain MBT76 (not shown).

Interestingly, while all verticillate *Streptomyces* also have orthologues of *ssgD* and *ssgG,* they lack an orthologue of *ssgE,* which is fully conserved in non-verticillate streptomycetes. Importantly, *ssgE* deletion mutants of *S. coelicolor* fail to produce long spore-chains, with instead single spores and occasional short spore chains formed, suggesting that SsgE plays a role in spore-chain length and morphogenesis (12). Verticillate streptomycetes produce short spore chains along the lateral wall of the aerial hyphae (Figure 1), and the lack of *ssgE* may directly correlate to this different mode of sporulation. Taken together, the absence of *ssgE* could be a useful taxonomic marker for verticillate streptomycetes, and it will be very interesting to see which other genetic differences may correlate to the sporulation phenotype seen in members of this clade.

### Chemotaxonomy

To further classify *Streptomyces* sp. MBT76 ^T^ and compare it to other streptomycetes, we performed chemotaxonomy based on chemical composition of the cell wall. A whole-cell hydrolysate of the strain was rich in LL-diaminopimelic acid, glucose, mannose and ribose. The predominant menaquinone (>25%) was MK9H8 (47%).The cellular fatty acids contained large proportions (>10%) of *anteiso*-C_15:0_ (34.40%), and *anteiso*-C_17:0_ (10.92%). Lower proportions (i.e. <10%) of *iso*-C_14:0_ (8.28%), *iso*-C_15:0_ (5.11%), iso-C_16:0_ (7.99%), *anteiso*-C_16:0_ (2.54%), C_16:1_ ω9 (2.84%), C_16:0_ (5.64%), C_18:1_ ω9 (8.93%), C_20:11_ ω11 (4.53%) and summed features C_18:2_ ω9,12/C_18:0_ (8.81%). Polar lipids are diphospatidylglycerol, phosphatidylethanolamine, phosphatidylinositol, Glycophosphatidylinositol, and an unknown lipid. Growth characteristics and phenotypic properties are summarized in Tables 1 and 2. Growth characteristics of *S. hiroshimensis* are summarized in Table S3.

### Description of *Streptomyces roseofaciens* sp. nov

*Streptomyces roseofaciens* (*ro.se.o.fa’ci.ens* L.adj.roseus.rosy; L.V.facie, make N.L.part.adj. roseofaciens, making rosy).

Aerobic, Gram-positive actinomycete which forms a branching substrate mycelium that carries aerial mycelium which forms verticillate chains with 3-5 smooth spores per chain. Substrate mycelium is pink on ISP1,2,3,4,5,7. Grows from 20 to 50°C, optimally at ~30°C, from pH 5.0 to pH 11, optimally at pH ~7, and in the presence of 2% NaCl. L-tyrosinase, hypoxanthine and casein are hydrolysed. Degrades starch and gelatine. Uses glucose, sucrose and inositol as sole carbon source. The strain is positive for acid and alkaline phosphatase, α-cysteine arylamidase, α-chymotrypsin, esterase (C4), esterase lipase (C8), α- and β-glucosidase, α-mannosidase, N-acetyl-β-glucosaminidase, napthol-AS-B1-phosphatase, trypsin and valine arylamidase, but negative for α-fucosidase, α- and β-galactosidase and β-glucoronidase (API-ZYM tests). The diagnostic amino acid in the peptidoglycan is LL-diaminopimelic acid, whole cell hydrolysates contain glucose, mannose and ribose. The predominant menaquinones is MK9(8). Polar lipids are diphospatidylglycerol, phosphatidylethanolamine, phosphatidylinositol, phosphatidylinositol mannosides, and an unknown lipid. The major fatty acids are anteiso-C_15:0_, and anteiso-C_17:0_. The strain has 44 putative secondary metabolite biosynthetic gene clusters predicted by antiSMASH 4.0 and has been analysed extensively by NMR- and MS-based metabolomics. The digital DNA G+C composition of the type strain is 71.9%.

The type strain (=NCCB 100637^T^ =DSM 106196^T^) was isolated from the QinLing mountains in the Shaanxi Province, China at an altitude of 660m, with permission. The species description is based on a single strain and hence serves as a description of the type strain. The GenBank accession number for the assembled genome of *Streptomyces roseofaciens* is GCA_001445655.1.

## Acknowledgements

This project was supported by an EMBO Short-Term Fellowship (6746) awarded to LvdA.

